# RWKV-IF: Efficient and Controllable RNA Inverse Folding via Attention-Free Language Modeling

**DOI:** 10.1101/2025.06.13.659654

**Authors:** Gaoyuan Ji, Kefan Xu, Chuhong Zheng

## Abstract

We present **RWKV-IF**, an efficient and controllable framework for RNA inverse folding based on the attention-free RWKV language model. By treating structure-to-sequence generation as a conditional language modeling task, RWKV-IF captures long-range dependencies with linear complexity. We introduce a decoding strategy that integrates Top-*k* sampling, temperature control, and G-C content biasing to generate sequences that are both structurally accurate and biophysically meaningful. To overcome limitations of existing datasets, we construct a large-scale synthetic training set from randomly generated sequences and demonstrate strong generalization to real-world RNA structures. Experimental results show that RWKV-IF significantly outperforms traditional search-based baselines, achieving higher accuracy and full match rate while greatly reducing edit distance. Our approach highlights the potential of lightweight generative models in RNA design under structural and biochemical constraints. The code is available at https://github.com/Lyttr/RWKVInverseFolding.git

## 1 Introduction

RNA, a molecule consisting of ribonucleotides in a chain-like structure, plays a pivotal role in the development of advanced therapeutics and biotechnologies. Designing RNA sequences that fold into specific secondary structures—known as the *RNA inverse folding* problem—is essential for enabling such applications. First introduced in the ViennaRNA package [14], RNA inverse folding involves generating RNA sequences that conform to a given target secondary structure.

However, this problem is computationally challenging. Traditional algorithms depended heavily on predefined structural configurations and domain-specific knowledge, restricting their adaptability for complex or novel design tasks [21, 7]. Moreover, RNA inverse folding is known to be NP-hard, which further limits the scalability of conventional approaches due to high computational costs [3].

Solving the RNA inverse folding problem has substantial practical significance. The ability to reliably generate functional RNA molecules with desired structural and functional properties enables a wide array of real-world applications. For example, in synthetic biology, engineered RNA molecules can act as biosensors or regulatory circuits [12, 5]. In therapeutics, structured RNA elements such as aptamers and small interfering RNAs (siRNAs) require precise folding to function effectively [16, 30].

In nucleic acid nanotechnology, rational RNA design underpins the construction of programmable RNA-based scaffolds and devices [13, 11, 1].

Historically, RNA design relied on empirical methods and directed evolution in wet-lab settings. These approaches were not only costly, but also time-consuming and inefficient, with a low success rate in identifying functional RNA [29].

To address the challenges, computational approaches are developed and rogressed from traditional search-based algorithms to modern learning-based approaches. Early search-based methods, such as RNAinverse [14], MCTS-RNA [33], and antaRNA [17], relied on heuristic optimization strategies to explore the sequence space. Later enhancements, including IncaRNAtion [25] and SAMFEO [35], integrated probabilistic sampling and ensemble-level insights for improved performance. More recently, deep learning has transformed the field, with models like SentRNA [28] and LEARNA[27] predicting sequences directly or using reinforcement learning, while advanced techniques such as Meta-LEARNA [26] and geometric deep learning models like RiboDiffusion [15] and RNAFlow [22] leverage meta-learning and structural priors to enhance generalization and capture RNA’s conformational diversity. Computational techniques offered an improvement, but still faced significant limitations, especially in generalization and efficiency for complex structures.

With the advent of deep learning, generative models—particularly large language models (LLMs)—have demonstrated impressive capabilities in natural language processing (NLP), as well as growing success in modeling biological and chemical sequences [20, 10]. While pre-trained language models have proven effective in predicting RNA-related properties [6, 2], their application to the generative side of RNA design, such as inverse folding, remains largely unexplored.

In this work, we explore the application of LLMs to the RNA inverse folding problem. Specifically, we investigate the use of RWKV, a lightweight, attention-free LLM architecture that combines the advantages of RNNs and Transformers.

In summary, our contributions are fourfold:

- We formulate RNA inverse folding as a conditional sequence generation problem and demonstrate the effectiveness of RWKV for modeling RNA structure-to-sequence mappings.
- We introduce a decoding strategy that combines Top-*k* sampling, temperature control, and logit-based G-C content adjustment to generate sequences that are both accurate and biophysically meaningful.
- We construct a large-scale synthetic training dataset by randomly generating RNA sequences. Despite being fully synthetic, we show that models trained on this dataset generalize well to real-world RNA structures, providing evidence that random sequence-based datasets are viable for training inverse folding models.
- We conduct extensive experiments on benchmark test sets, demonstrating that our approach significantly outperforms traditional search-based baselines in both accuracy and structural fidelity.

## 2 Related Work

### 2.1 Concept and Objectives of RNA Inverse Folding

As shown in Figure 1, RNA inverse folding refers to the problem of designing RNA sequences that can fold into a given target RNA structure (typically secondary or tertiary structure). As a core challenge in RNA design, its objectives include not only finding sequences that match the target structure but also ensuring their functional activity in biological environments and meeting specific constraints (e.g., sequence length, GC content). This problem holds significant importance in synthetic biology, drug development, and other fields.

**Figure 1.**
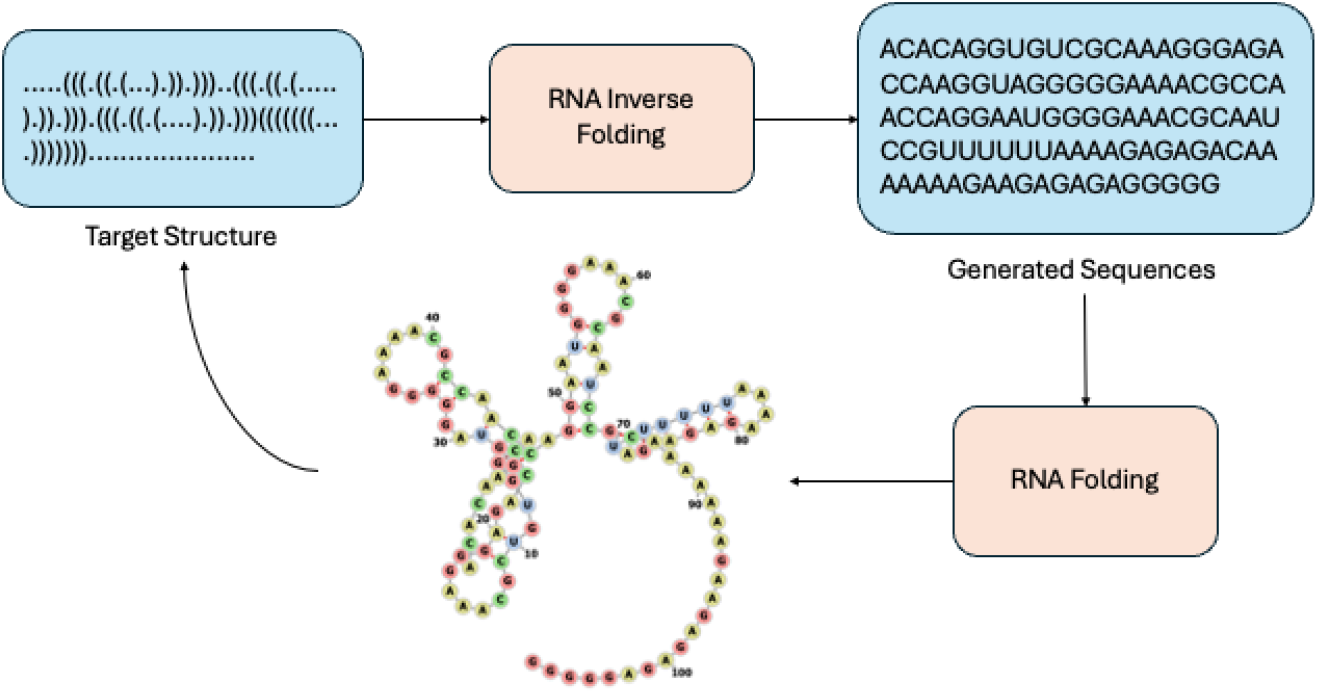
RNA Inverse Folding Pipeline

### 2.2 Search-based Methods

These approaches systematically explore the sequence space to identify RNA sequences that fold into the target structure. Early methods such as **RNAinverse** [14, 19, 4], which is among the earliest algorithms, employs an adaptive random walk strategy starting from a randomly initialized sequence. It iteratively mutates nucleotides to minimize the base pair distance between predicted and target secondary structures. **MCTS-RNA** [33] utilized Monte Carlo tree search to explore candidate sequences by simulating and evaluating the folding properties of different sequences. Additionally, **antaRNA** [17] applied ant colony optimization, where sequences are generated via weighted random sampling. The search weights are updated based on the quality of previously generated sequences, progressively refining the sequence pool. **IncaRNAtion** [25] focuses on generating sequences that not only fold into the target structure but also match specified GC content using Boltzmann sampling with stochastic backtracking. **SAMFEO** [35] integrates both structure-level and ensemble-level information to guide iterative sampling, mutation, and optimization, generating a large number of viable RNA sequences.

Search-based methods suffer from several key limitations. Firstly, they often rely on simplified energy models or heuristic scoring functions, making it challenging to accurately capture the complexity of RNA folding, particularly tertiary structure interactions. Secondly, these methods exhibit poor computational efficiency when dealing with long sequences or complex structures, as the search space grows exponentially with sequence length. Moreover, local search strategies are prone to getting trapped in suboptimal solutions and fail to explore the entire sequence space, resulting in limited sequence diversity.

### 2.3 Learn-based Methods

With the development of deep learning, learn-based approaches have become the mainstream in RNA inverse folding. Early methods such as **SentRNA** [28] employs a fully connected neural network trained end-to-end using human-designed RNA sequences, directly predicting sequence assignments for a given structure. **LEARNA** [26] employed deep reinforcement learning frameworks to generate sequences that fold into target secondary structures through interaction with an environment. After that, **Meta-LEARNA** [26] extends LEARNA using meta-learning techniques to improve generalization across diverse RNA structures, treating each structure as a distinct Markov Decision Process (MDP). Furthur more, **Meta-LEARNA-Adapt** [26] combines Meta-LEARNA’s pre-trained policy with fine-tuning using LEARNA to adapt to new tasks.

Recent advances in geometric deep learning, such as **RiboDiffusion** [15] and **RNAFlow** [22], have shown promising results. RiboDiffusion introduced a generative diffusion model that learns the sequence distribution conditioned on fixed backbone structures, iteratively transforming random initial sequences into candidates that satisfy the target tertiary structure. RNAFlow combined flow matching with pre-trained structure prediction networks and considered multiple RNA conformations to enhance the modeling of RNA dynamics.

Despite significant progress, learn-based methods still face several challenges. Current models exhibit limited performance when dealing with complex RNA structures (e.g., those containing pseudoknots or multi-chain interactions) and are highly dependent on the quality and quantity of training data. Many approaches consider only a single optimal structure during prediction, neglecting the conformational flexibility of RNA, which is a key determinant of its function. Additionally, most models lack direct constraints on sequence functionality during design, resulting in sequences that may be structurally correct but lack biological activity.

## 3 Method

We propose to use the RWKV model (specifically the Eagle/RWKV-5 architecture [24]) to perform RNA inverse folding as a conditional sequence generation task. Given a target secondary structure in dot-bracket notation, the model generates an RNA sequence expected to fold into that structure.

The input follows the format as <Structure>\n<Sequence>. Each pair is tokenized using a character-level tokenizer, where the dot-bracket structural notation and corresponding nucleotide sequences are mapped to discrete token indices. The input structural tokens are processed through a stack of cascaded RWKV blocks, each comprising a time-mix subblock and a channel-mix subblock. This architecture enables efficient modeling of long-range dependencies in RNA structures. The final hidden state is passed through a prediction head to predict the next nucleotide token in an autoregressive manner.

The RWKV model architecture is characterized by four key components in its time-mixing and channel-mixing blocks:

- **R** (Receptance): Controls how much historical information flows through the network.
- **W** (Weight): Learnable position-weighting parameters that decay with relative distance.
- **K** (Key): Operates similarly to the key in standard attention mechanisms for content retrieval.
- **V** (Value): Functions like the value in conventional attention, storing features for aggregation.

These core components interact at each timestep through multiplicative operations, as illustrated in Figure 2. The information flows sequentially through each processing stage within the block.

**Figure 2.**
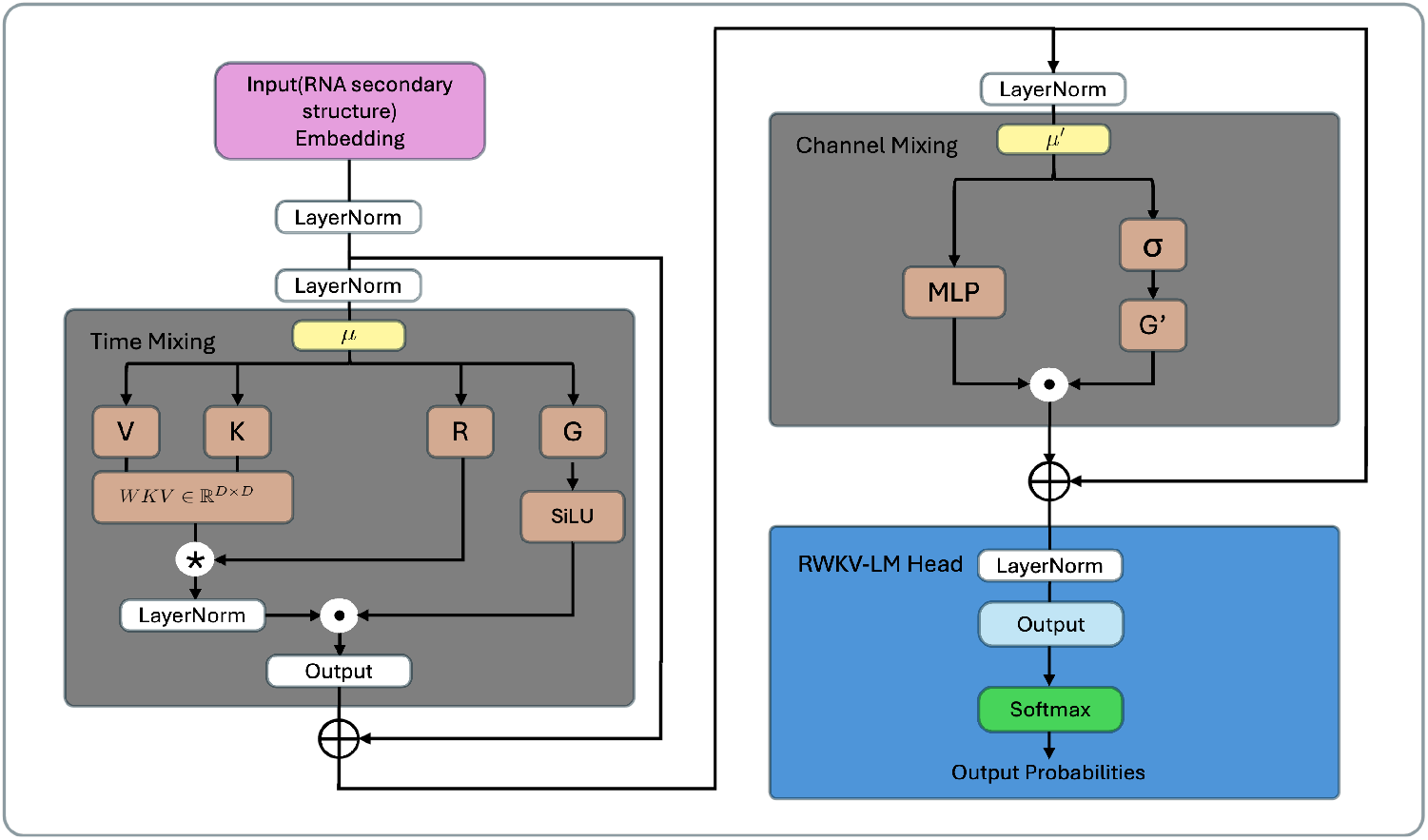
RWKV architecture overview

The RWKV architecture processes RNA secondary structure embeddings through a series of identical residual blocks, each performing sequential transformations in two distinct computational stages before producing outputs for the RWKV-LM Head.

As illustrated in Figure 2, each residual block first applies Time Mixing to capture position-aware temporal patterns across the sequence, followed by Channel Mixing that enables nonlinear interactions between feature channels. This dual-stage design allows the model to efficiently learn both sequential dependencies and cross-feature relationships within a unified recurrent framework.

### 3.1 Time-Mixing Block

The Time Mixing sublayer extends RWKV-4’s formulation through several key modifications while preserving its efficient recurrent structure. The core computation follows these transformations:

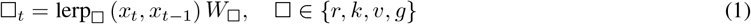

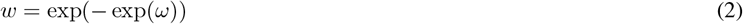

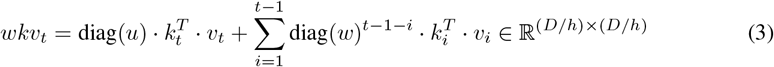

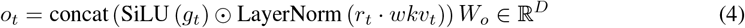

where *D* is the model dimension and *h* is the number of heads.

The computation begins by projecting the interpolated inputs (*x*_*t*_ and *x*_*t*−1_) through learned weights *W*_□_ to obtain the receptance (*r*_*t*_), key (*k*_*t*_), value (*v*_*t*_), and gate (*g*_*t*_) terms. The attention mechanism employs a novel decay formulation where *w* = exp(− exp(*ω*)) ensures contractive properties, with *ω* ∈ ℝ*D/h* as trainable head-specific parameters. The *wkv*_*t*_ term combines both immediate context (through the *u*-boosted current token contribution) and historical information (via the decaying sum of past 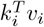terms).

The LayerNorm operation applies group-wise normalization across the *h* attention heads, equivalent to Group-Norm [32] with *h* groups. The final output *o*_*t*_ integrates these components through a gated composition: the normalized attention outputs are modulated by SiLU-activated gates before projection through *W*_*o*_.

The attention computation admits an efficient recurrent implementation:

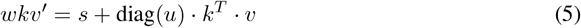

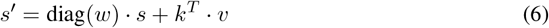

This formulation reveals Eagle’s connection to linear attention mechanisms, where the *wkv* term serves as a decay-based analogue to normalized key-value products. Each channel in the recurrent state *s* maintains a decaying accumulation of past *k*^*T*^ *v* terms, with channel-specific decay rates determined by *w*. The current token receives special treatment through the learned boost *u* before combining with the historical state. The receptance vector *r*_*t*_ then acts similarly to queries in linear attention, selecting relevant information from this dynamically maintained memory.

### 3.2 Channel-Mixing Block

The Eagle architecture retains the Channel Mixing module from RWKV-4 with minimal modifications, primarily reducing the hidden dimension from 4*D* to 3.5*D* to maintain parameter parity after introducing new gating mechanisms in the Time Mixing sublayer. The computation follows the same fundamental operations as RWKV-4, beginning with linear interpolation of the current and previous inputs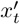 and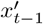to generate the receptance and key vectors:

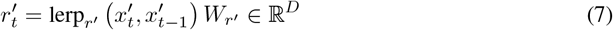

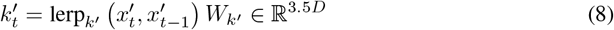

The value computation then applies a ReLU activation followed by squaring before projection through *W*_*v*_*′*, creating a non-linear feature transformation:

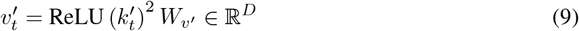

The final output combines these components through sigmoid-gated modulation, where the receptance vector 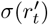 controls information flow from the transformed features 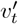:

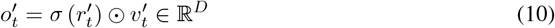

### 3.3 Token Shift

Eagle incorporates the Token Shift mechanism from previous RWKV architectures, implementing a lightweight temporal mixing operation mathematically equivalent to a 1D causal convolution with kernel size 2. The core operation employs learnable linear interpolation between consecutive tokens:

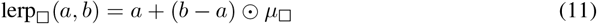

where each interpolation operation (for □ ∈ {*r, k, v, g*}) uses a dedicated learnable mixing vector

*µ*_□_ ∈ ℝ*D*. This design enables three fundamental capabilities:

- **Per-channel temporal blending**: Each dimension in the receptance (*r*), key (*k*), value (*v*), and gate (*g*) vectors autonomously determines its optimal balance between current (*x*_*t*_) and previous (*x*_*t*−1_) token information
- **Head-specific mixing patterns**: Independent *µ*_□_ parameters per attention head facilitate specialized temporal processing strategies
- **Induction head formation**: The mechanism directly enables the emergence of induction heads [9] within single layers, as individual heads can maintain separate subspaces for current and historical token processing

### 3.4 Top-*k* Sampling and G-C Content Control

To generate RNA sequences in a controlled yet diverse manner, we adopt the Top-*k* sampling strategy during decoding. At each time step *t*, given the input token sequence *x*_1:*t*_, the model outputs a logit vector *z*_*t*_ over the vocabulary. These logits are first scaled by a temperature parameter *τ* to control randomness:

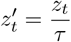

To further guide sequence generation toward a desired guanine-cytosine (G-C) content ratio, we introduce a content-aware adjustment to the logits. Let *r*_*t*_ denote the current G-C ratio in the generated sequence *x*_1:*t*−1_:

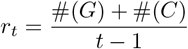

With a target G-C ratio *r*_target_, a bias *δ* is applied to the logits of tokens corresponding to G and C bases based on the deviation from this target:

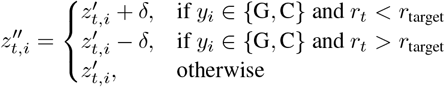

This adjustment encourages the model to prefer G or C tokens when the current ratio is below the target, and suppresses them when the ratio is above.

After that, the softmax function is added to the adjusted logits:

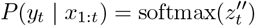

Instead of sampling from the full distribution, we restrict sampling to the top *k* most probable tokens:

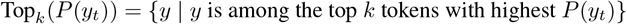

The probabilities of the top *k* tokens are then renormalized:

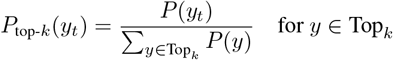

Finally, the next token *y*_*t*_ is sampled from this reduced distribution:

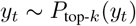

This approach balances stochastic sampling and biochemical constraints, allowing us to generate RNA sequences that are both diverse and biophysically meaningful.

## 4 Dataset Building

Since datasets contribute significantly in the training of LLMs, the datasets for LLM-based RNA inverse folding are expected to meet the following requirements:

- **Large scale structure labels**: Without any predefined knowledge and rule, the datasets bring all information of RNA structures that the model can learn from. Although unlabeled RNA sequences can be used for RNA feature extraction tasks, the structure-sequence pairs are necessary for inverse folding. Therefore, the scale of labeled RNA datasets directly affects the performance of the model.
- **Strong structure and sequence diversity**: The generalizability of the model are strictly limited by the diversity of datasets, including structure-level diversity and sequence-level diversity. The former enables the model to maintain the performance on new structures, while the latter ensures the model can learn the deep pattern of RNAs, rather than simply maps one sequence with one structure.
- **Low cost folding evaluation**: Since RNA inverse folding is a generative task, one input structure is mapped with multiple output RNA sequences. Therefore, an evaluation tool is required to verify the effectivity of output. Specifically, an RNA folding tool is used to obtain the structure of output sequence and compare it with the input.

However, existing datasets suffer from the lack of both structure labels and diversity, along with the absence of evaluation methods. For instance, the Eternabench-cm dataset [31]contains only 12293 samples with 2199 unique structures, and the top 10% most frequent structures cover 74% samples. Similarly, nearly half of the RNAs in bpRNA-1m [8] dataset are repeated. Moreover, since the structures in bpRNA-1m are obatined by experiments, computational RNA folding methods, such as RNAfold, do not restore the structures from the sequences, which prevents the evaluation.

To address the problem, we built a dataset with 2000000 randomly generated RNA sequences. In details, 2082 unique structures from Eternabench-cm dataset are used as the test set after filtering extremely long samples. After that, sequences in the training set, with length between 80 and 120, are obtained by randomly selecting nucleotides (A, U, G, C). Then, the RNAfold from ViennaRNA package is adopted to generate their structures in the form of dot-bracket notations. There is no intersection between training set and test set.

Table 1 provides the statistics information of the training set. The randomly generated training set and Eternabench-cm test set share the similar statistics features, suggesting that models trained on the random sequences are expected to work on RNA datasets in real world. Moreover, with the higher average max depth and max span, the structures in the training set are more challenging than structures in the test set.

**Table 1:** Statistics of RNA secondary structure datasets.

| Statistic | Test Set | Training Set |
| --- | --- | --- |
| Average base pairs | 24.44 | 27.79 |
| Average max depth | 13.96 | 19.25 |
| Average max span | 45.44 | 69.68 |
| Average structure length | 105.55 | 102.51 |
| Total structures | 2,082 | 2,000,000 |
| Unique structures | 2,082 | 2,000,000 |

## 5 Experiment Settings

Since the training strategy of vanilla RWKV is designed for language generation tasks, it is adapted for the RNA inverse folding task. The original approach flattens the training tokens into a large 1-dimension vector and randomly samples a window from it. However, it will destroy the structure-sequence pairs. Therefore, we use the data with the shape of <Pairs, Tokens> and sampling by rows, enabling the model to learn relations from complete structure-sequence pairs. Moreover, a loss mask is added to the structure part, helping the model focus on predicting the sequences.

An RWKV model with 8 layers, an embedding size of 512, a context length of 256, and a vocabulary size of 9 was deployed. The learning rate schedule employed linear warm-up followed by cosine annealing, starting from an initial learning rate of 1e-4 and decaying to a final learning rate of 1e-5. The model was trained for 7 epochs, during which the loss converged to 0.5. Training was performed on an NVIDIA A5000 GPU.

For the evaluation, the trained model generated one sequence per unique structure in the test set. After that, the RNAfold was used to obtain the structures of these sequences, which were compared with the input structures. Three metrics were selected: average character-level accuracy, average edit distance and full match accuracy. The RNAinverse tool from ViennaRNA package and AntaRNA served as the baseline.

## 6 Result

According to Table 2, the RWKV-IF provides better performance than baselines on all metrics. While differences on character-level accuracy is not obvious, a huge gap can be observed on the edit distance and full match accuracy, which reduce by 83.4% and improve by 89.2%, respectively. It proves that although all approaches can generate sequences with structures similar with the target, our method performs much better on obtaining a perfectly matched sequence. In other words, our approach requires fewer attempts to generate an expected sequence, reducing the computational cost of large scale generation tasks.

**Table 2:**
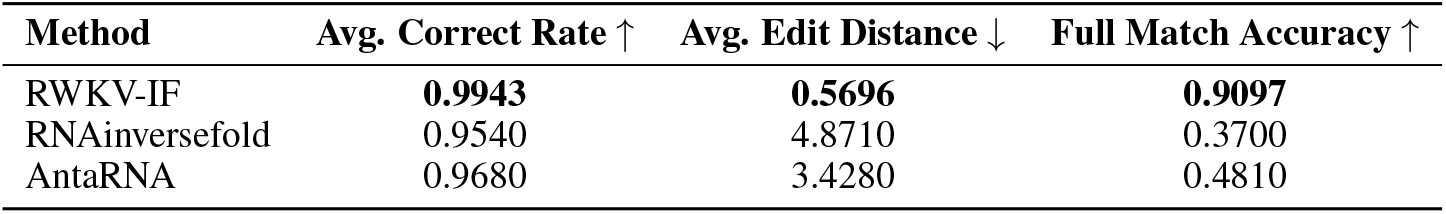
Performance comparison on the RNA inverse folding task.

Table 3 reports the performance of RNA sequence generation under different temperature values using Top-*k* sampling. As the temperature decreases, the decoder becomes more deterministic, leading to higher average correctness and full match accuracy along with lower distance among sequences. Note that with the temperature lower than 0.01, the performance remains undamaged and average distance achieves around 15. The result proves the model’s effectiveness of both accuracy and diversity. Table 4 summarizes the results of G-C content control of RNA sequences. The actual G-C ratios closely match the specified target values, demonstrating that the proposed logit adjustment method effectively steers the nucleotide composition of generated sequences. The temperature is set to 0.01 in this experiment.

**Table 3:**
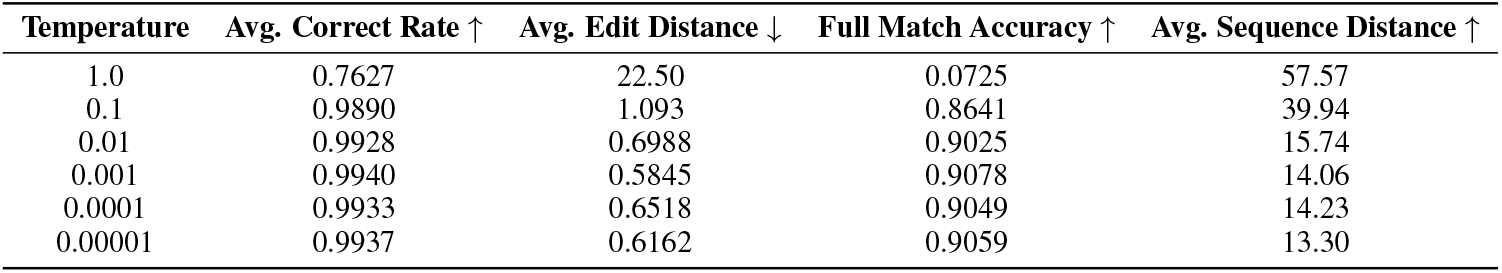
RNA Diversity with Top-*k* Sampling.

| Temperature | Avg. Correct Rate ↑ | Avg. Edit Distance ↓ | Full Match Accuracy ↑ | Avg. Sequence Distance ↑ |
| --- | --- | --- | --- | --- |
| 1.0 | 0.7627 | 22.50 | 0.0725 | 57.57 |
| 0.1 | 0.9890 | 1.093 | 0.8641 | 39.94 |
| 0.01 | 0.9928 | 0.6988 | 0.9025 | 15.74 |
| 0.001 | 0.9940 | 0.5845 | 0.9078 | 14.06 |
| 0.0001 | 0.9933 | 0.6518 | 0.9049 | 14.23 |
| 0.00001 | 0.9937 | 0.6162 | 0.9059 | 13.30 |

**Table 4:** G-C Content Control Results.

| Target GC | Actual GC | Avg. Correct Rate ↑ | Avg. Edit Distance ↓ | Full Match Accuracy ↑ |
| --- | --- | --- | --- | --- |
| 0.2500 | 0.2545 | 0.9151 | 8.5793 | 0.5115 |
| 0.5000 | 0.4972 | 0.9731 | 2.6849 | 0.7824 |
| 0.7500 | 0.7457 | 0.9549 | 4.4083 | 0.7205 |

## 7 Discussion

### 2.1 Advantage of RWKV

Inspired by [34], RWKV was proposed as an attention-free, RNN-like architecture for NLP tasks [23]. In contrast to Transformer-based approaches, RWKV offers several distinctive advantages, including efficient handling of long-range dependencies, sequential token modeling, and linear time and memory complexity, making RWKV particularly well-suited for the RNA inverse folding task, where capturing global structural dependencies over long structure representations is critical. Moreover, RNA inverse folding often involves repeating sampling and optimization under structural and thermodynamic constraints. RWKV’s lightweight inference and streaming capabilities achieve efficient generation of candidate sequences with acceptable computational costs especially for large scale generation and huge sequence length.

### 7.2 Future Work

In future work, we plan to incorporate additional benchmark datasets such as Eterna100v2 dataset [18] to further evaluate model generalization across diverse structural distributions and performance on challenging structure requirements. Expanding the set of baseline methods will also provide a more comprehensive comparison against recent state-of-the-art approaches. Moreover, we aim to extend our framework to support RNA structures with pseudoknots, which are difficult for existing approaches and critical for accurately capturing functional RNA conformations.

## 8 Conclusion

In this work, we propose RWKV-IF, a novel LLM-based approach to the RNA inverse folding problem. By framing inverse folding as a conditional sequence generation task, our method effectively models long-range dependencies within RNA structures with high computational efficiency. Through the integration of Top-*k* sampling, temperature control, and G-C content control mechanism, our model achieves high accuracy,structural diversity, and controllable nucleotide composition. Extensive experiments demonstrate that RWKV-IF outperforms traditional search-based methods, reducing edit distance by 83.4%, and improving full match accuracy by 89.2% comparing with selected baselines. These results highlight the potential of generative language models in advancing computational RNA design and open up new directions for scalable and flexible biomolecular sequence generation.

## References

[1] Kirill A Afonin, Mathias Viard, Alexey Y Koyfman, Angelica N Martins, Wojciech K Kasprzak, Martin Panigaj, Ravi Desai, Arti Santhanam, Wade W Grabow, Luc Jaeger, et al. Multifunctional rna nanoparticles. Nano letters, 14(10):5662–5671, 2014.

[2] Manato Akiyama and Yasubumi Sakakibara. Informative rna base embedding for rna structural alignment and clustering by deep representation learning. NAR genomics and bioinformatics, 4(1):lqac012, 2022.

[3] Jeff Anderson-Lee, Eli Fisker, Vineet Kosaraju, Michelle Wu, Justin Kong, Jeehyung Lee, Minjae Lee, Mathew Zada, Adrien Treuille, Rhiju Das, et al. Principles for predicting rna secondary structure design difficulty. Journal of molecular biology, 428(5):748–757, 2016.

[4] Mirela Andronescu, Zhi John Zhang, and Anne Condon. Efficient parameter estimation for rna secondary structure prediction. Bioinformatics, 21(16):3504–3511, 2004.

[5] James Chappell, Melissa K Takahashi, and Julius B Lucks. Creating small transcription activating rnas. Nature chemical biology, 11(3):214–220, 2015.

[6] Jiayang Chen, Zhihang Hu, Siqi Sun, Qingxiong Tan, Yixuan Wang, Qinze Yu, Licheng Zong, Liang Hong, Jin Xiao, Tao Shen, et al. Interpretable rna foundation model from unannotated data for highly accurate rna structure and function predictions. arXiv preprint 2204.00300, 2022.

[7] Alexander Churkin, Matan Drory Retwitzer, Vladimir Reinharz, Yann Ponty, Jérôme Waldispühl, and Danny Barash. Design of rnas: comparing programs for inverse rna folding. Briefings in bioinformatics, 19(2):350–358, 2018.

[8] Padideh Danaee, Mason Rouches, Michelle Wiley, Dezhong Deng, Liang Huang, and David Hendrix. bprna: large-scale automated annotation and analysis of rna secondary structure. Nucleic acids research, 46(11):5381–5394, 2018.

[9] Nelson Elhage, Neel Nanda, Catherine Olsson, Tom Henighan, Nicholas Joseph, Ben Mann, Amanda Askell, Yuntao Bai, Anna Chen, Tom Conerly, Nova DasSarma, Dawn Drain, Deep Ganguli, Zac Hatfield-Dodds, Danny Hernandez, Andy Jones, Jackson Kernion, Liane Lovitt, Kamal Ndousse, Dario Amodei, Tom Brown, Jack Clark, Jared Kaplan, Sam McCandlish, and Chris Olah. A mathematical framework for transformer circuits. Transformer Circuits Thread, 2021. https://transformer-circuits.pub/2021/framework/index.html.

[10] Daniel Flam-Shepherd, Kevin Zhu, and Alán Aspuru-Guzik. Language models can learn complex molecular distributions. Nature Communications, 13(1):3293, 2022.

[11] Wade W Grabow and Luc Jaeger. Rna self-assembly and rna nanotechnology. Accounts of chemical research, 47(6):1871–1880, 2014.

[12] Alexander A Green, Pamela A Silver, James J Collins, and Peng Yin. Toehold switches: de-novo-designed regulators of gene expression. Cell, 159(4):925–939, 2014.

[13] Peixuan Guo. The emerging field of rna nanotechnology. Nature nanotechnology, 5(12):833– 842, 2010.

[14] Ivo L Hofacker, Walter Fontana, Peter F Stadler, L Sebastian Bonhoeffer, Manfred Tacker, Peter Schuster, et al. Fast folding and comparison of rna secondary structures. Monatshefte fur chemie, 125:167–167, 1994.

[15] Wenhao Huang, Haotian Shi, D. Wu, et al. Ribodiffusion: Generative diffusion model for rna backbone and sequence co-design. In International Conference on Learning Representations (ICLR), 2024.

[16] Anthony D Keefe, Supriya Pai, and Andrew Ellington. Aptamers as therapeutics. Nature reviews Drug discovery, 9(7):537–550, 2010.

[17] Rick Kleinkauf, Matthias Mann, Rolf Backofen, and Sebastian Will. antarna: ant colony-based rna sequence design. In Proceedings of the 2015 ACM Conference on Bioinformatics, Computational Biology, and Health Informatics, pages 219–226. ACM, 2015.

[18] Rohan V. Koodli, Boris Rudolfs, Jonathan Romano, Hannah K. Wayment-Steele, William A. Dunlap, Eterna Participants, and Rhiju Das. Redesigning the eterna100 for the vienna 2 folding engine. bioRxiv, 2025.

[19] Ronny Lorenz, Stephan H Bernhart, Christian Honer zu Siederdissen, Hakim Tafer, Christoph Flamm, Peter F Stadler, and Ivo L Hofacker. Rnainverse 2.0: Improvements and extensions to the rnainverse rna secondary structure design algorithm. Nucleic Acids Research, 39(Suppl_2):W494–W498, 2011.

[20] Mehrsa Mardikoraem, Zirui Wang, Nathaniel Pascual, and Daniel Woldring. Generative models for protein sequence modeling: recent advances and future directions. Briefings in Bioinformatics, 24(6):bbad358, 2023.

[21] David H Mathews, Walter N Moss, and Douglas H Turner. Folding and finding rna secondary structure. Cold Spring Harbor perspectives in biology, 2(12):a003665, 2010.

[22] Niveditha Nori, Shirui Liu, Linyi Zhang, et al. Rnaflow: Generative flow matching for rna backbone and sequence co-design. In International Conference on Learning Representations (ICLR), 2024.

[23] Bo Peng, Eric Alcaide, Quentin Anthony, Alon Albalak, Samuel Arcadinho, Stella Biderman, Huanqi Cao, Xin Cheng, Michael Chung, Matteo Grella, Kranthi Kiran GV, Xuzheng He, Haowen Hou, Jiaju Lin, Przemyslaw Kazienko, Jan Kocon, Jiaming Kong, Bartlomiej Koptyra, Hayden Lau, Krishna Sri Ipsit Mantri, Ferdinand Mom, Atsushi Saito, Guangyu Song, Xiangru Tang, Bolun Wang, Johan S. Wind, Stanislaw Wozniak, Ruichong Zhang, Zhenyuan Zhang, Qihang Zhao, Peng Zhou, Qinghua Zhou, Jian Zhu, and Rui-Jie Zhu. Rwkv: Reinventing rnns for the transformer era, 2023.

[24] Bo Peng, Daniel Goldstein, Quentin Anthony, Alon Albalak, Eric Alcaide, Stella Biderman, Eugene Cheah, Teddy Ferdinan, Haowen Hou, Przemysław Kazienko, et al. Eagle and finch: Rwkv with matrix-valued states and dynamic recurrence. arXiv preprint 2404.05892, 3, 2024.

[25] Vladimir Reinharz, Yann Ponty, and Jérôme Waldispühl. Incarnation: efficient generation, storage and sampling of rna sequences with fixed desired properties. Bioinformatics, 29(24):3082– 3088, 2013.

[26] Fabian Runge, Lars Lorch, Sven Schneider, Annika Hess, Kai Böhm, Katharina Bieker, Christian Kling, Stefan Pfister, Michael Hiller, and Björn Ronacher. Learna: Reinforcement learning for rna design. In International Conference on Learning Representations (ICLR), 2019.

[27] Frederic Runge, Danny Stoll, Stefan Falkner, and Frank Hutter. Learning to design rna, 2019.

[28] Jinbo Shi, Rhiju Das, et al. Sentrna: Improving rna design by integrating a human-derived prior with deep learning. Scientific Reports, 8(1):1–12, 2018.

[29] Craig Tuerk and Larry Gold. Systematic evolution of ligands by exponential enrichment: Rna ligands to bacteriophage t4 dna polymerase. science, 249(4968):505–510, 1990.

[30] Jie Wang, Ze Lu, M Guillaume Wientjes, and Jessie L-S Au. Delivery of sirna therapeutics: barriers and carriers. The AAPS journal, 12:492–503, 2010.

[31] Hannah K Wayment-Steele, Wipapat Kladwang, Alexandra I Strom, Jeehyung Lee, Adrien Treuille, Alex Becka, Eterna Participants, and Rhiju Das. Rna secondary structure packages evaluated and improved by high-throughput experiments. Nature methods, 19(10):1234–1242, 2022.

[32] Yuxin Wu and Kaiming He. Group normalization. In Proceedings of the European conference on computer vision (ECCV), pages 3–19, 2018.

[33] Ye Yang, Jian Li, et al. Rna design via deep reinforcement learning. In Advances in Neural Information Processing Systems, volume 30, 2017.

[34] Shuangfei Zhai, Walter Talbott, Nitish Srivastava, Chen Huang, Hanlin Goh, Ruixiang Zhang, and Josh Susskind. An attention free transformer, 2021.

[35] Jiahua Zhou, Lianhui Zhang, Wen Wang, et al. Samfeo: Structure and ensemble aware multifaceted rna design optimization. Nature Communications, 14(1):4311, 2023.

